# Microbial cysteine degradation is a source of hydrogen sulfide in oxic freshwater lakes

**DOI:** 10.1101/2021.11.30.467465

**Authors:** Patricia Q. Tran, Samantha C. Bachand, Jacob C. Hotvedt, Kristopher Kieft, Elizabeth A. McDaniel, Katherine D. McMahon, Karthik Anantharaman

## Abstract

The sulfur-containing amino acid cysteine is abundant in the environment including in freshwater lakes. Biological cysteine degradation can result in hydrogen sulfide (H_2_S), a toxic and ecologically relevant compound that is a central player in biogeochemical cycling in aquatic environments. Here, we investigated the ecological significance of cysteine in oxic freshwater lake environments, using isolated cultures, controlled growth experiments, and multi-omics. We screened bacterial isolates enriched from natural lake water for their ability to produce H_2_S when provided cysteine. In total, we identified 29 isolates that produced H_2_S and belonged to the phyla *Bacteroidetes, Proteobacteria,* and *Actinobacteria*. To understand the genomic and genetic basis for cysteine degradation and H_2_S production, we further characterized 3 freshwater isolates using whole-genome sequencing (using a combination of short-read and long-read sequencing), and quantitatively tracked cysteine and H_2_S levels over their growth ranges: *Stenotrophomonas maltophilia* (Gammaproteobacteria), *Stenotrophomonas bentonitica* (Gammaproteobacteria) and *Chryseobacterium piscium* (Bacteroidetes). We observed a decrease in cysteine and increase in H_2_S, and identified genes involved in cysteine degradation in all 3 genomes. Finally, to assess the presence of these organisms and genes in the environment, we surveyed a five-year time series of metagenomic data from the same isolation source (freshwater Lake Mendota, WI, USA) and identified their presence throughout the time series. Overall, our study shows that sulfur-containing amino acids can drive microbial H_2_S production in oxic environments. Future considerations of sulfur cycling and biogeochemistry in oxic environments should account for H_2_S accumulation from degradation of organosulfur compounds.

**Importance:** Hydrogen sulfide (H_2_S), a naturally occurring gas with biological origins, can be toxic to living organisms. In aquatic environments, H_2_S production typically originates from anoxic (lacking oxygen) environments such as sediments, or the bottom layers of thermally stratified lakes. However, the degradation of sulfur-containing amino acids such as cysteine, which all cells and life forms rely on, can be a source of ammonia and H_2_S in the environment. Unlike other approaches for biological H_2_S production such as dissimilatory sulfate reduction, cysteine degradation can occur in the presence of oxygen. Yet, little is known about how cysteine degradation influences sulfur availability and cycling in freshwater lakes. In our study, we identified diverse bacteria from a freshwater lake that can produce H_2_S in the presence of O_2_. Our study highlights the ecological importance of oxic H_2_S production in natural ecosystems and necessitates a change in our outlook of sulfur biogeochemistry.

## Introduction

In most natural environments, hydrogen sulfide gas (H_2_S) production is usually attributed to defined groups of bacteria and archaea (1, 2), and occurs primarily in anoxic environments. During the process of dissimilatory sulfate reduction, sulfate acts as a terminal electron acceptor, and is converted to hydrogen sulfide. However, other pathways for H_2_S production exist, namely assimilatory sulfate reduction, in which H_2_S contributes to cell growth and increased biomass, and the desulfurylation (desulfurization) of sulfur-containing amino acids such as cysteine which can lead to production of pyruvate, ammonia, and H_2_S (3). It is believed that assimilatory sulfate reduction contributes to growth but does not release H_2_S from the cell, while dissimilatory sulfate reduction and cysteine degradation can both contribute to growth and release ecologically relevant nitrogen and sulfur compounds into the ecosystem.

The sulfur cycle is composed of several assimilatory and dissimilatory pathways, which interact in complex ways through biotic and abiotic factors. Sulfur cycling in freshwater ecosystems can be ecologically significant, especially in places where strong redox gradients exist (4). For example, in high arctic lakes, sulfur-compounds are suggested to serve as biogeochemical hubs (5). Cysteine, a sulfur containing amino acid, is proposed to be an overlooked source of both carbon (6) and sulfur. Additionally, seston (particles in water comprised of both living and non-living organisms) contains organosulfur containing lipids which settle into the sediments, and contributes to the sulfur pool, even in highly oligotrophic lakes such as Lake Superior (7).

In seasonally stratified lakes consisting of oxygenated warm water (epilimnion) floating atop colder anoxic waters (hypolimnion), H_2_S is often abundant in the hypolimnion (8, 9), due to oxygen demand driving terminal electron acceptor depletion. However, an overlooked player in the pool of available H_2_S are the use of organosulfur compounds, such as cysteine, by microbes. Cysteine is required to produce many proteins and is also important for protein structure. It is one of the two amino acids (methionine being the other) that contains a sulfur group; however, the sulfhydryl group on cysteine is more reactive and can lead to H_2_S formation. Like all amino acids, cysteine also contains an amine group that will form ammonia once the molecule is degraded. As such, cysteine degradation (desulfurylation) by microbes leads to H_2_S production. H_2_S is ecologically relevant because it can be toxic to plants and animals. During periods of anoxia, H_2_S can accumulate to levels beyond the threshold for living organisms, and can cause massive fish kills (10). Unlike other H_2_S sources, cysteine desulfurylation can occur under oxic conditions, thereby expanding the environmental scope of this sulfur pool. Indeed, cysteine can be desulfurylated under oxic conditions in the laboratory, but little information exists on the natural prevalence of this process in lakes. We expect that H_2_S production in oxic environments (during the mixed water column periods of the year, and throughout the stratified period in the mixed epilimnion) could result from cysteine breakdown by microbes.

In this study, we investigated the prevalence of organosulfur degradation (desulfurylation) in a freshwater lake, using both laboratory and genomic evidence, to advance our understanding of oxic sulfur cycling in aquatic ecosystems (**Figure 1**). First, we grew bacterial isolates enriched from Lake Mendota’s oxic epilimnion to quantify H_2_S and ammonia production, which informs the potential for organosulfur degradation in an oxygenated aquatic environment. We found 18 isolates producing H_2_S under oxygenic conditions. We selected three H_2_S-producing isolates for detailed characterization using full-genome sequencing and chemical analyses to track cysteine concentrations and H_2_S accumulation during their growth: *Stenotrophomonas maltophilia* (Gammaproteobacteria), *Stenotrophomonas S. bentonitica* (Gammaproteobacteria) and *Chryseobacterium piscium* (Bacteroidetes). In all three isolates, cysteine decreased and H_2_S increased over their exponential growth curve under oxic conditions. Finally, we contextualized our laboratory results using a time-series of metagenomic data from the same isolation source (Lake Mendota, WI, USA), to study the temporal importance of organosulfur degradation. We found that genes for cysteine desulfurylation were present and abundant throughout the time-series suggesting that the ability to degrade cysteine is well represented in Lake Mendota.

**Figure 1.**
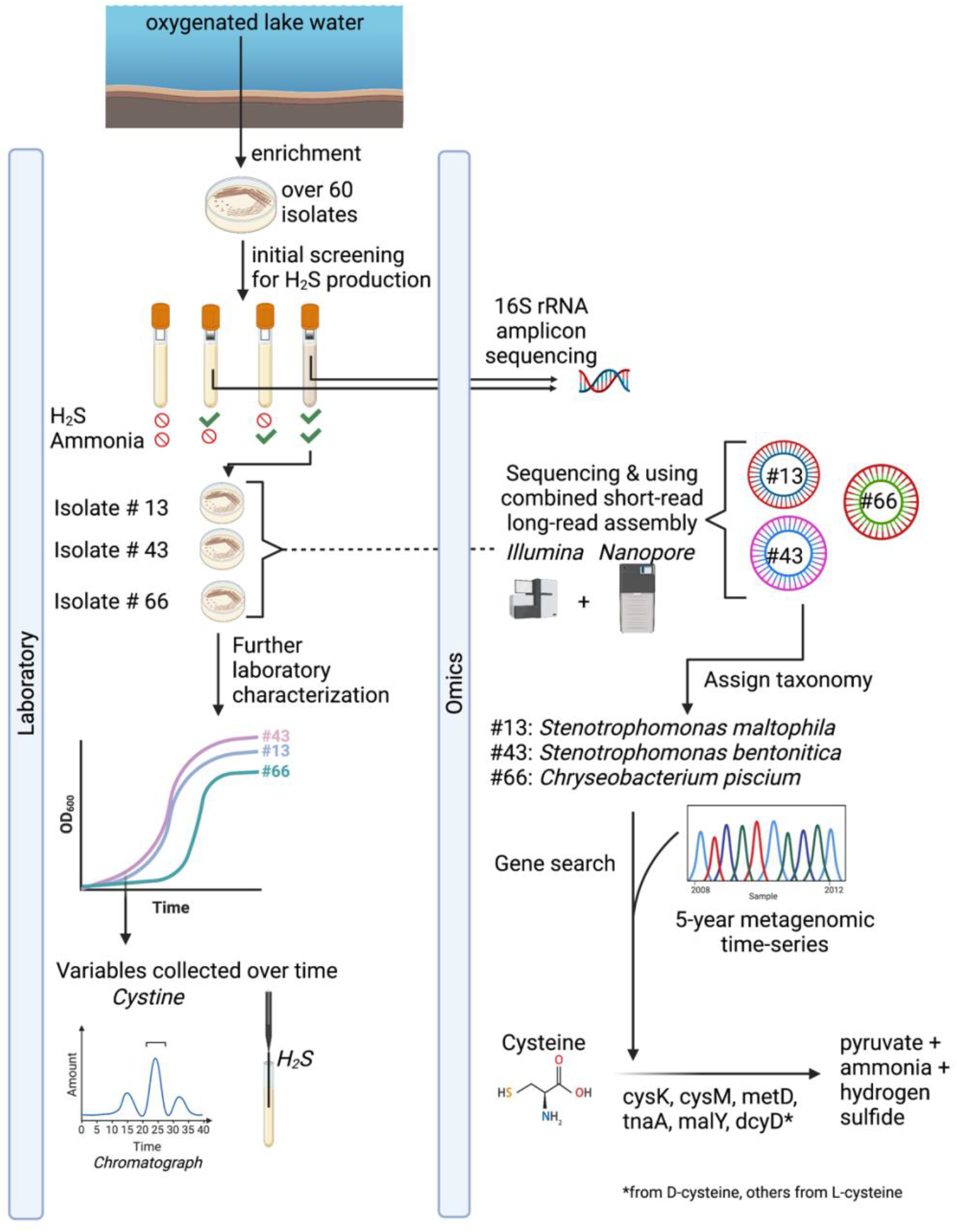
Overview of methods used in this study. Isolates were enriched from oxygenated lake water. Isolates were screened for H_2_S and ammonia production, and those that produced H_2_S were selected for 16S rRNA sequencing. Then based on the results, three isolates were selected for whole-genome sequencing using a combination of short and long-reads. Genome characterization of functional potential and taxonomic classification was conducted on the isolates. Screening of genes involved in cysteine degradation was conducted in the isolates, and a five-year metagenomic time series of Lake Mendota (2008-2012).

## Results

### Isolates capable of H_2_S production in oxic conditions

To answer the question of whether bacteria could produce H_2_S in the presence of oxygen, we grew pure culture isolates originally recovered from the water column of temperate eutrophic Lake Mendota. We grew the isolates in moderately rich media (R2A, see methods) under control and treatment conditions (cysteine addition) and tracked H_2_S production after 24 hours (**Figure S1, Table S1**). Using qualitative H_2_S measurements, we found that 18 isolates produced H_2_S and ammonia when grown in the presence of amended cysteine. We performed 16S rRNA gene sequencing on the 29 isolates that produced H_2_S, regardless of whether they produced ammonia or not when amended with cysteine. Isolates that produced both H_2_S and ammonia were identified as *Stenotrophomonas rhizophila* (Betaproteobacteria), *Stenotrophomonas maltophilia* (Betaproteobacteria), *Citrobacteria gillenii* (Gammaproteobacteria), and *Chryseobacterium sp.* (Bacteroidetes), whereas those producing H_2_S but not ammonia were identified as *Pseudomonas arsenicoxydans, Pseudomonas mandelii, Pseudomonas migulae, Pseudomonas thivervalensis,* and *Microbacterium flavescens*.

### Detailed microbiological, chemical and genomic characterization in selected isolates

Next, we selected 3 isolates (#43, #13 and #66) representing distinct species based on 16S rRNA sequence taxonomy (97% identity), that produced H_2_S for further characterization. These detailed characterizations include OD600-based growth rates and paired quantitative measurements of cysteine and H_2_S concentrations in the spent medium. Cysteine addition resulted in concommital H_2_S production over time (**Figure 2A**, **Table S2**). All organisms used L-cysteine preferably to D-cysteine (**Figure 2B**, **Table S3**).

**Figure 2.**
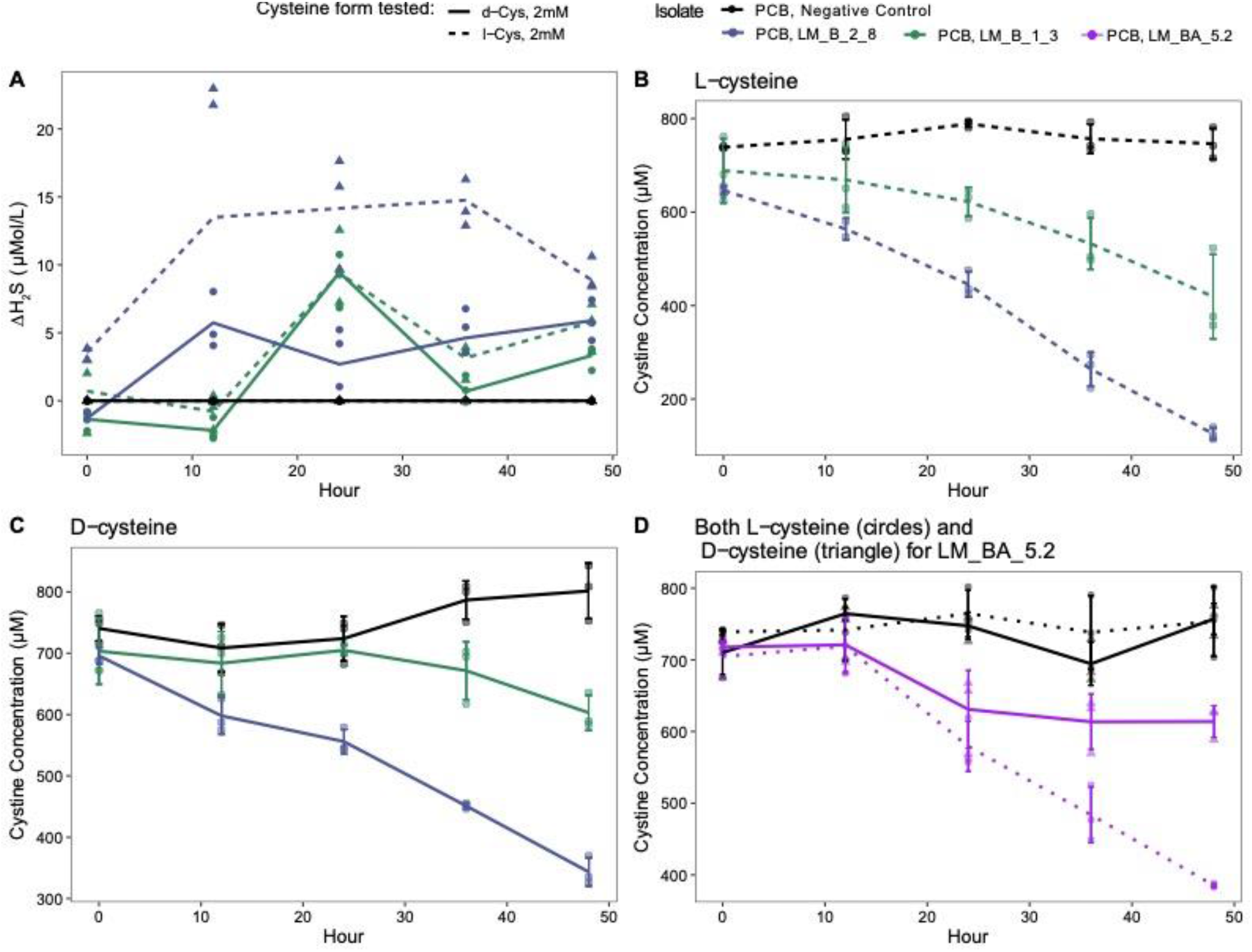
Further characterization of the three isolates and demonstration of cysteine degradation and H_2_S production. A. Higher amounts of H_2_S was produced by the three isolates over 50 hours compared to negative controls. B, C, D. Identification of different forms of cysteine that can be degraded. L-cysteine decreased in all isolates compared to the control (B, D). D-cysteine also decreased over time in all samples except the negative control, however, the net amount decreased was less compared to L-cysteine. Cysteine concentrations were measured as cystine as described in the methods. Because isolates #13 and #43 were assessed in a different experimental run than #66, but using the exact same instruments and methods, plots B, C and D are separated by HPLC runs. Due to large sample volume it was not possible to test all isolates and conditions in one HPLC run.

Next, we performed whole-genome sequencing using combined short-read and long-read sequencing on these 3 isolates (**Table 1**). We performed whole-genome sequencing because functional information such as gene content cannot be predicted reliably from 16S rRNA sequencing alone. The full genome of Isolate 43 was assembled into a single circular genome, and taxonomically assigned to *Stenotrophomonas maltophilia*. Unlike the 16S rRNA sequence which assigned it to *S. rhizophila*, the full-genome was actually closer to *S. maltophilia*. The full genome of isolate #13, could be assembled into 2 long contigs, and was taxonomically assigned to *S. bentonitica*. The *Chryseobacter* genome was assembled into one circular genome, and assigned to *Chryseobacterium piscium*. All three genomes were estimated to be 100% complete based on CheckM. Overall, the 16S rRNA gene amplicon sequencing performed prior agreed with full-genome sequencing assignment in some cases, and in others, the whole-genome sequencing assignment allowed finer taxonomic resolution (such as in the case of Isolate 13). Overall, whole-genome sequencing provided more information about the isolates’ metabolic potential. The genomic content was then used to inform how or why H_2_S might be produced in oxic environments, as shown in the laboratory experiments.

**Table 1.** Genome characteristics of 3 selected isolates that produce H_2_S in the presence of oxygen and cysteine.

| Name | Genome Size (bp) | Completeness<sup>1</sup> | GC content | Taxonomy<sup>2</sup> | Number of contigs |
| --- | --- | --- | --- | --- | --- |
| 13-LM-B-02-08 (referred to as #13) | 4,188,104 | 100% | 66.8% | d__Bacteria;p__Proteobacteria;c__Gammaproteobacteria;o__Xanthomonadales;f__Xanthomonadaceae;g__Stenotrophomonas;s__**Stenotrophomonas maltophilia_O** | 1 |
| 43-LM-B-01-03 (referred to as #43) | 4,325,715 | 100% | 66.5% | d__Bacteria;p__Proteobacteria;c__Gammaproteobacteria;o__Xanthomonadales;f__Xanthomonadaceae;g__Stenotrophomonas;s__**Stenotrophomonas bentonitica** | 2 |
| LM_BA_5.2 (referred to as #66) | 1,375,102 | 100% | 33.7% | d__Bacteria;p__Bacteroidota;c__Bacteroidia;o__Flavobacteriales;f__Weeksellaceae;g__Chryseobacterium;s__**Chryseobacterium piscium** | 7 |

<sup>1</sup> Using CheckM (See Methods) <sup>2</sup> Using GTDB-tk (See Methods)

We used gene annotations of the 3 isolates to infer the presence of genes involved in cysteine metabolism. We identified genes involved in cysteine degradation to ammonia, pyruvate, and H_2_S: *metC*, *malY*, *tnaA*, *cysM*, *cysK* (which involve the use of L-cysteine as the substrate), and *dcyD* (which involve the use of D-cysteine as the substrate) (**Table S4**). However, we note that these genes may have other enzymatic activities, such as cysteine biosynthesis instead of degradation (**Table S5**). (**Figure 3**)

**Figure 3.**
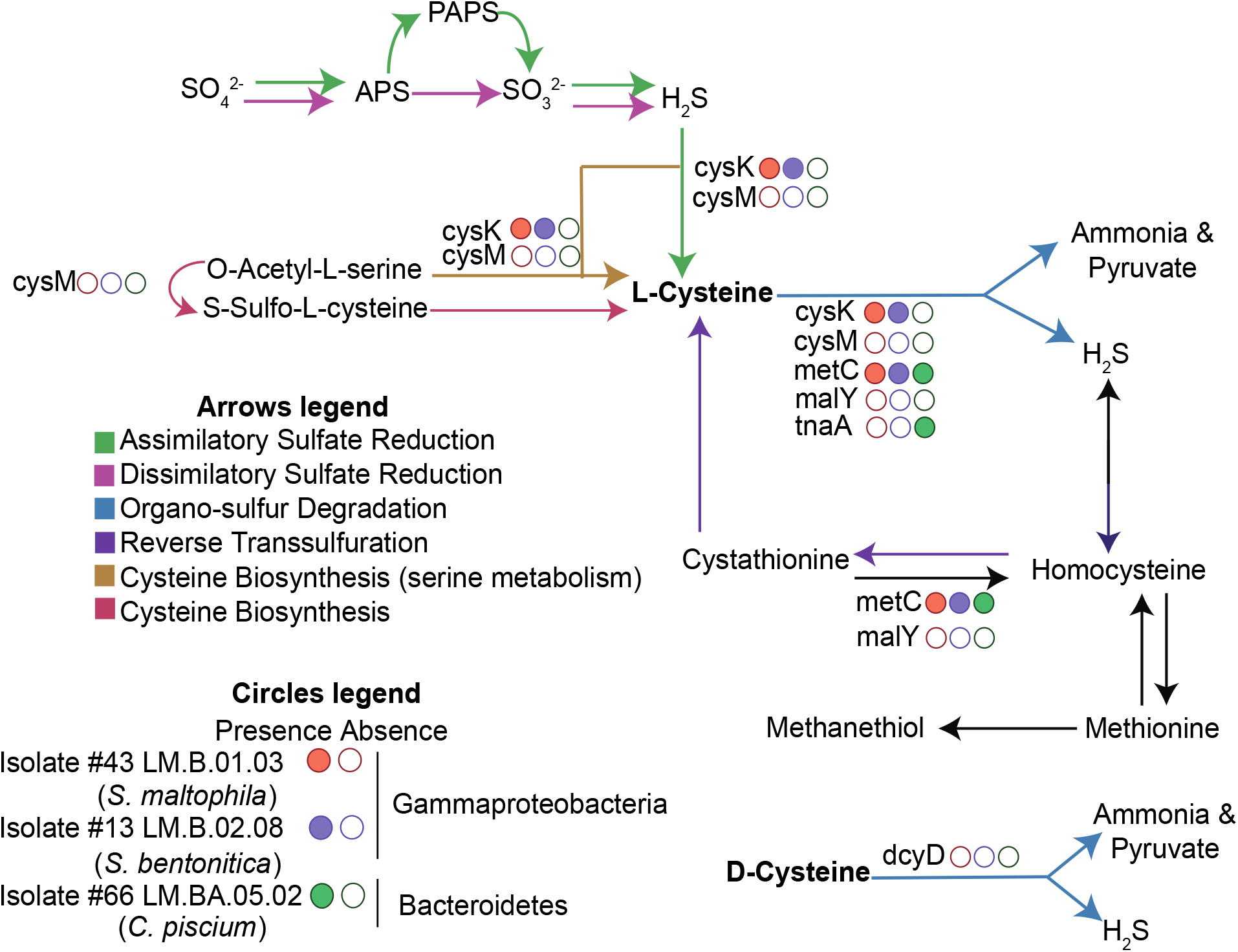
Diagram showing pathways for sulfur and organosulfur metabolism. Several pathways for hydrogen sulfide and cysteine production exist in microorganisms. The presence/absence of key genes (*cysK*, *cysM*, *malY*, *metC*, *tnaA*, *sseA*, *aspB*, and *dcyD* along the blue arrows) in the three isolate genomes are shown by filled (present) circles, and unfilled (not present) circles.

Leveraging the full-genomic content of the 3 isolates (**Table S6**), we proposed a joint cellular map based on identified metabolic functions and pathways in the genomes (**Figure 4A**). All three isolates contained pathways associated with central carbon metabolism: including the TCA cycle, glycolysis, gluconeogenesis, the pentose phosphate pathway and the glyoxylate cycle. They could all generate fatty acids using fatty acid biosynthesis, and catabolize fatty acids through the beta-oxidation pathway. As expected, they had genes for cysteine metabolism including *metC*, *malY*, *tnaA*, *cysM*, *cysK*, and *dcyD*, cysteine biosynthesis pathways from homocysteine and serine, as well pathways for degradation of other amino acids including methionine.

Despite these similarities, the three isolates also have distinguishing characteristics amongst them (**Table S6,** **Figure 4B**). For example, while all isolates encoded genes for sulfur oxidation (sulfur dioxygenase), genes for thiosulfate oxidation were present in the two *Stenotrophomonas* isolates but not *Chryseobacterium*. The *Chryseobacterium* isolate encoded for a urease suggesting the use of organic nitrogen in the form of urea but this was absent in the two *Stenotrophomonas* isolates. Finally, genes for sugar utilization were identified in the two *Stenotrophomonas* isolates but not in *Chryseobacterium*.

**Figure 4.**
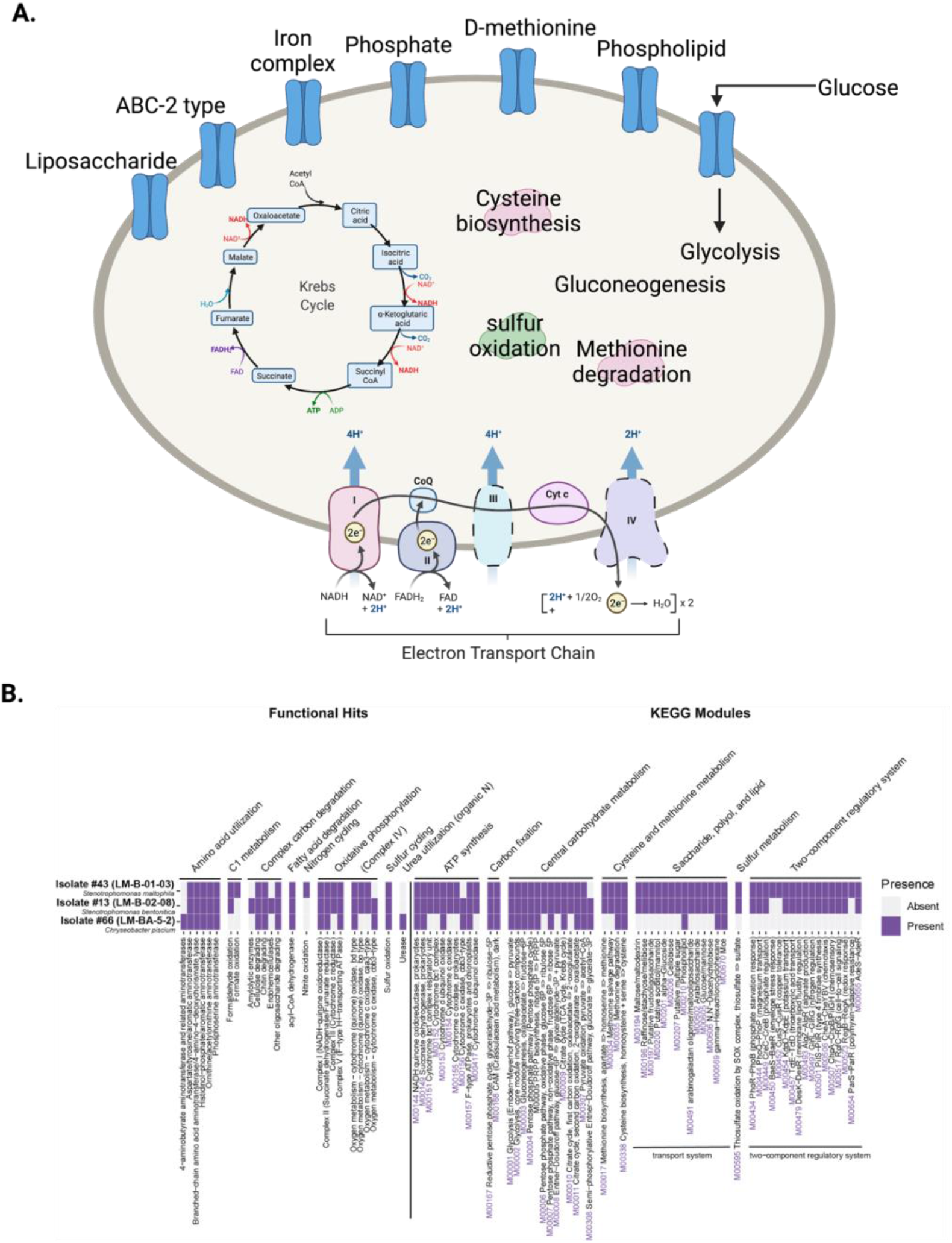
A. Cellular map showing important metabolic pathways and transporters which were common to all three isolates. A complete list is found in **Table S6**. The KEGG module identifiers are listed in purple whenever relevant. **B. Heat map showing selected metabolic functions and pathways in the three isolates.** A complete list is found in **Table S6**.

### Presence of cysteine-degrading organisms and genes in a five-year metagenomic environmental time-series

To put these laboratory results and lab-grown organisms into a natural environment context, we leveraged a previously published metagenomic time-series collected from the oxygenated upper mixed layer of Lake Mendota spanning 2008-2012 (97 time points) to search for the presence of cysteine degradation genes in the microbial communities.

First, we searched the time-series to see if organisms in our study were also present in the time series. To do this, we linked the 16S rRNA gene sequences of the isolated organisms to the assembled metagenomes (i.e. contigs) from the time series. We found that while the 16S rRNA sequences were also present in the time series (**Table S7, S8, S9**), and broadly distributed over time, these scaffolds were not part of binned genomes. Therefore, little information about these isolates would be gathered from metagenomic data only. As such, the full-genome sequencing we performed was particularly helpful in understanding the full genomic structure of the H_2_S-producing organisms.

Second, we searched for the 6 genes associated with cysteine degradation and H_2_S production (**Table S5**) (in binned and unbinned contigs) using KEGG HMMs with hmmsearch. In total, we searched over 22 million amino acid sequences and identified 1882 hits to the 5 marker genes found in the isolate metagenomes, *dcyD* was not found (**Figure 5**, **Table S10**). *cysK* and *malY* were the genes with the most corresponding matches at any time point, followed by *metC* and *cysM*. Only 2 scaffolds contained *tnaA*. Overall, after correcting for genome size, there was no obvious temporal trend of the genes, although genes were found throughout the 5-year time series.

Among these cysteine-degrading gene sequences, several were identified in binned metagenome-assembled genomes (MAGs) (**Figure 5B**, **Table S11**), which allowed for the assignment of taxonomy. Overall, we identified 139 genes to be distributed in genomes of organisms from Actinobacteria, Bacteroidetes, Chloroflexota, Cyanobacteria, Planctomycetes, Proteobacteria, and Verrucomicrobia, representing common freshwater lineages (11). The *tnaA* gene was only present in Bacteroidetes, but other genes were more broadly distributed among taxonomic groups.

**Figure 5.**
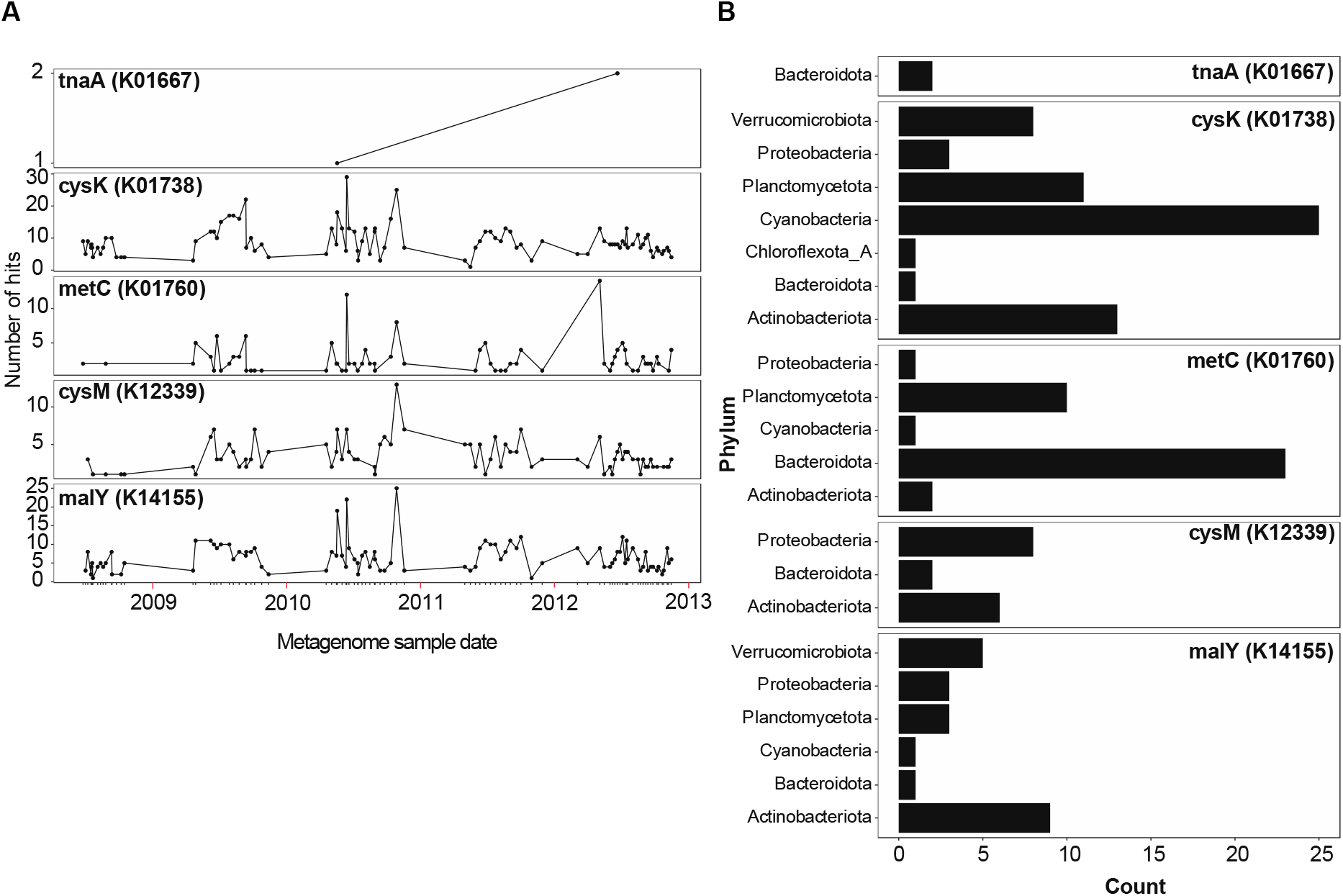
A. Counts of cysteine desulfurylation genes (5 genes) in a 5-year time-series of Lake Mendota. We also searched for dcyD (6th gene) but did not identify it in any sample. **B. Presence of these 5 genes in metagenome-assembled metagenomes from the same time series.**

## Discussion

### Different types of H_2_S production: the fate of cysteine and the origin of the H_2_S

The H_2_S-producing isolates fell into two groups when grown in media with cysteine: ammonia-producing and ammonia-consuming. We hypothesize that those able to produce ammonia and H_2_S in the presence of cysteine, but not under “control” conditions were those that were potentially contributing to the H_2_S pool in the lake. However, it is possible that the isolates that consume ammonia also conduct the cysteine degradation pathway, but do not excrete ammonia from the cell. Instead, since ammonia is an important metabolic precursor, it could be used by the organism instead of being released into the media, making it undetectable with our analytical technique.

We do not believe that this occurs with H_2_S, and that most, if not all, of the isolates that produce H_2_S in the presence of oxygen will excrete it from the cell. H_2_S is toxic to organisms that undergo aerobic cellular respiration (12). One potential reason to keep H_2_S in the cell is that some bacteria appear to use H_2_S as a protective compound against antibiotics, but H_2_S accumulation also creates a large amount of stress on the cell making it useful only in extreme situations (13). While it is possible that organisms may use H_2_S internally as a sulfur source, we did not identify any genes encoding sulfide quinone oxidoreductases, flavocytochrome c dehydrogenases, or other genes for the oxidation or transformation of H_2_S. All three isolates were obligate aerobes based on laboratory assays.

Of the 20 amino acids, little is known about cysteine in the environment. One of the difficulties in studying the fate of cysteine in oxic environments is that it can be abiotically and spontaneously oxidized into cystine (14), which *Escherichia coli* has been shown to uptake (15). In a study of *E.coli* K-12 that lacked a cysteine transporter, cysteine could enter the cell through transporters dedicated to other amino acids, when no amino acids alternatives other than cysteine were present in the medium (16). It is likely that the majority of H_2_S produced by the cell in the three isolates originates from cysteine given the demonstrated reduction in cystine concentrations when the isolates are grown with added cysteine (**Figure 2**), coupled with the accumulation of H_2_S and ammonia.

### Genomic structure of the H_2_S-producing isolates

Overall, the three isolates selected for whole genome sequencing revealed genes for cysteine degradation into H_2_S. Based on laboratory studies, they were able to produce H_2_S in the presence of oxygen. The isolates were obligate aerobes, which presents interesting questions about the life history of these organisms.

Little information is known about the ecology of *Stenotrophomonas maltophilia*, *Stenotrophomonas bentonitica*, and *Chryseobacterium piscium* in the natural environment. *S. maltophilia* is a cosmopolitan bacterium in nature, and found in a range of natural environments, particularly in association with plants(17). *S. bentonitica* was originally characterized in bentonite formations, was predicted to have high tolerance to heavy metals (18), and has been observed in arctic seawater (19). *C. piscium* was isolated from a fish in the arctic ocean (20), but its ecological significance in the oceans remains unknown. This previously described *C. piscium* strain LMG 23089 was not reported to produce H_2_S yet our genetic analyses suggest that it has the enzymatic machinery to degrade cysteine.

One possible explanation for this discrepancy is that LMG 23089 was previously grown on SIM medium to test H_2_S, which is lower resolution than the modern H_2_S probes which measure µM concentrations, and because SIM medium uses thiosulfate as a sulfur source. As a side test on isolate #66, H_2_S was not produced when thiosulfate was provided, but H_2_S was produced when cysteine was provided.

One particular finding of this study was that none of the 6 genes searched for cysteine degradation into H_2_S and ammonia was common to *all* three isolates, despite all 3 isolates showing the same cysteine-decrease, ammonia-increase, and H_2_S-increase over time. This could be explained by alternative, perhaps less straightforward pathways for H_2_S production. One pathway is led by a gene named cystathionine gamma-lyase (“CTH” or “CSE”). In some bacteria and mammals, this enzyme is involved in H_2_S production (21). An HMM search for this enzyme showed that it was present in Isolate #13, 43 and #66. While it was not initially included in the initial methods and study, this could hint to another commonality among oxic H_2_S producing organisms.

### Challenges associated with measuring oxic H_2_S production from organosulfur in the environment

Extrapolating these laboratory results to widespread distribution of organosulfur degradation in the natural environment necessitates several steps, namely because of the major knowledge gaps that exist concerning the sulfur cycle in freshwater lakes, and because bridging the gap between cultivation-based, omics-based (22), and field-based experiments is needed. Foremost, the identity, distribution, and availability of organosulfur compounds broadly across lakes globally is currently mostly unknown. Cysteine is notoriously difficult to measure, and many previous studies characterizing the amino acid composition of the water column only measure the sulfur-containing organosulfur compound taurine (23, 24).

Organic sulfur in the form of cysteine is an important organosulfur amino acid, and is important in protein folding and function (25). As such, there is a difference in the fates of cysteine when it exists bound in cell walls, versus when cysteine is free in the water column and available for degradation by bacteria. While cysteine has been shown to contribute to the carbon pool and carbon flow in lakes (6), more quantitative field measurements are necessary to support whether cysteine also serves as a sulfur pool. Yet, other forms of organosulfur have important significance in aquatic environments. In marine environments, for example, DMSP (dimethylsulfoxonium propionate) is a critical component of the marine organosulfur cycle (26).

Additionally, current differences between computational gene similarity searches versus *in vivo* enzymatic functions are challenging to assess for the genes responsible for the cysteine degradation into pyruvate, ammonia, and H_2_S. One reason is that the enzymatic activity of the gene has mostly been described in model organisms such as *E. coli*, and it has been shown that gene expression can be induced by genetic factors, or environmental factors such as metals (27). At least 6 genes have been proposed to have this enzymatic activity, yet each gene may serve different functions *in situ*, and it is difficult to assert directionality of enzymatic function based on metagenomic or genomic analyses only. To this end, the isolated bacterial strains from this study, which are non-model organisms, and originate from the natural freshwater lake environment, may be used for further detailed biochemical, physiological, and microbiological studies. Further characterization of these bacterial isolates using gene knockout, gene induction, or heterologous gene expression studies may inform the functional activity of these genes in nature.

### Implications of oxic H_2_S production by microbes in freshwaters

This study demonstrates that H_2_S production by microbes in lake ecosystems occurs in the presence of oxygen, using evidence from pure culture bacterial isolates, and screening of long-term metagenomic time-series. By combining lake-to-laboratory experiments, we show that multiple bacterial strains spanning Gammaproteobacteria, Betaproteobacteria, Actinobacteria, and Bacteroidetes are all able to produce H_2_S under oxic conditions, and at temperatures that would be ecologically relevant for surface lake water during the summer. Surface water temperatures in Lake Mendota can reach up to 27°C, and the top few meters of water surface are saturated in oxygen. Worldwide, maximum lake surface temperature can range between 23 to 31°C (28).

Unlike dissimilatory sulfate reduction, cysteine use by bacteria to generate ammonia, pyruvate, and H_2_S, is not dependent on sulfate as an initial reaction substrate. Increased sulfate concentrations are shown to lead to higher sulfate reduction rates in shallow eutrophic freshwater, the sulfur originating from algal decay for example (29). While Lake Mendota is a low-iron and high-sulfate lake (30), not all lakes have elevated sulfate levels, and therefore, H_2_S production might previously not have been thought of as relevant to study. However, sulfur-containing amino-acids can have many origins. In lakes, concentrations of amino acids (free dissolved and combined) often reflect the input and outputs of the lake (31, 32). For example, amino acids contributed a detectable amount to the nitrogen cycle, and bacterial utilization of amino acids contributes to nitrogen pool and cycling in natural ecosystems, (32) although cysteine amino acids were not measured in that study.

Freshwater lakes that are dimictic can become stratified in temperature and oxygen during the summer, and oxygen concentrations vary throughout the year. In the fall and spring, oxygen is abundant, and cysteine degradation into H_2_S could be a relevant process for the sulfur pool, and H_2_S fluxes to the atmosphere could be significant since wind is prevalent. Under ice during the winter, where oxygen is plentiful, H_2_S could be produced but could be consumed or oxidized. On the other hand, gases would be trapped under ice. During summer, the anoxic hypolimnion and sediments are known H_2_S sources due to dissimilatory sulfate reduction, but density gradients would prevent H_2_S from reaching the atmosphere. However, the oxygenated mixed epilimnion could be an H_2_S source through organosulfur degradation. If we consider the importance of oxic hydrogen sulfide production, which could occur year-round, the H_2_S pool and the scope of sulfur transformations may be greater than anticipated, if we focus solely on the anoxic hypolimnion (**Figure 6**). Future work aiming to understand the broader distribution of sulfur-containing amino acids and other organosulfur compounds in freshwaters, their fates and transformations, as well as their contribution to H_2_S production, will inform global sulfur biogeochemical cycling.

**Figure 6.**
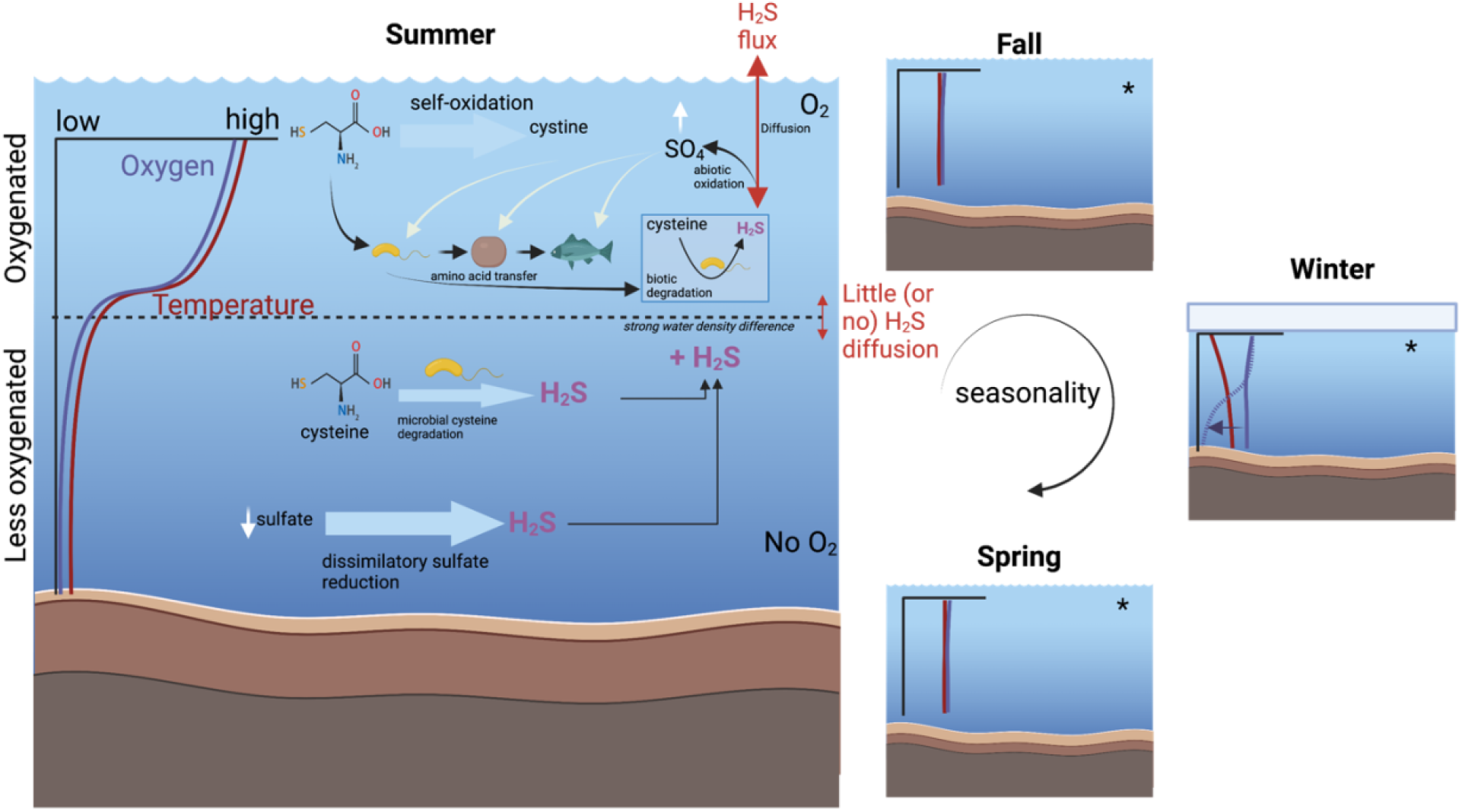
Conceptual figure showing the potential relevance of cysteine degradation in a freshwater lake environment, with respect to oxygen availability and seasonality. Oxygenated seasons and part of the lake water columns are shown with an asterisk. Significant research gaps include cysteine concentrations in the natural environment over time, hypothesized H_2_S fluxes across different layers in the lake water column, and contribution of different H_2_S sources in the hypolimnion. In all season, portions of the water column can be oxygenated.

## Methods

### Enrichment cultures of isolates from a temperate freshwater lake

Lake Mendota (43°06′24″ N 89°25′29″ W) is a temperate eutrophic lake in South-Central Wisconsin, in Madison, WI, USA. Lake Mendota is part of the Long-Term Ecological Research Network North Temperate Lakes (NTL-LTER, https://lter.limnology.wisc.edu/about/lakes). Lake Mendota encounters annual stratification and annual seasonal anoxia in the hypolimnion. Lake water was collected on September 14, 2018 from an integrated water sample (0m to 12m) from Lake Mendota at the “Deep Hole” station (43°05’54”, 89°24’28”), where the maximum depth is 23.5m. The water samples were collected during stratification from the oxygenated epilimnion. The lake water was collected in pre-acid washed 2 L sampling bottles using a flexible PVC tube, and brought back on shore within hours for immediate processing. Serial dilution was performed and lake bacteria were grown on PCB (plate count media broth) agar media, at room temperature (∼21°C), in the lab in the light. The PCB media was made of: 1 L water, 5g/L of yeast extract, 10g/L of tryptone and 2g/L of dextrose/D-glucose. If grown on solid media, 10g of agar per 1L media was added. Enrichment resulted in about 60 isolates.

### Screening for cysteine degradation into H_2_S and ammonia

Isolates were able to grow on PCB and R2A media. R2A media is a culture medium for bacteria that typically grow in freshwater. It is less “nutrient rich” than PCB media, and therefore slightly closer to natural lake water. For the screening of the isolates for H_2_S production, we grew them on R2A media. Each isolate had two treatments: grown in R2A media without cysteine for the control, and grown in R2A media with amended cysteine as the treatment.

R2A media consisted of (1 L of water): 0.5g of casein, 0.5g of dextrose, 0.5g of starch, 0.5g of yeast extract, 0.3g of K2HPO4, 0.3g of sodium pyruvate, 0.25g of peptone, 0.25g of beef extract, 0.024g of MgSO4, and autoclave. To make the same media for plates, we added 15g of agar before autoclaving. For controls, isolates were grown in the media without cysteine amendments. For “treatments”, 2mM cysteine was added.

To assess the amount of cysteine degradation into H_2_S and ammonia, we screened each of the 60 isolates for H_2_S and/or ammonia accumulation. To test H_2_S production, we grew the strains individually in liquid media, and used lead acetate test strips (Fisher Scientific, USA) to qualitatively assess H_2_S accumulation in the headspace. A darkening of the strip shows that H_2_S was produced. To test ammonia concentrations after 24 hours, we measured samples at time zero and 24 hours using Ammonia Salicylate Reagent Powder Pillows and Ammonia Cyanurate Reagent Powder Pillows (Hatch Reagents) and used spectrophotometry at the 655nm wavelength.

### Identification of H_2_S producing bacteria using 16S rRNA sequencing

Colony PCR and DNA extractions were conducted using the EtNa Crude DNA Extraction and ExoSAP-ITTM PCR Product Cleanup protocols on the isolates that tested positive for producing H_2_S (10). Full length 16S rRNA gene products were generated for sequencing using universal 16S rRNA primers (27f, 1492r)(33).DNA concentration yields were measured using the QuBit dsDNA HS assay kit (QuBit). DNA was sequenced at the University of Wisconsin-Madison Biotechnology Center (Madison, WI, USA) The program 4Peaks (34) was used to clean the base pairs by quality-checking followed by homology search using BLASTn against the NCBI Genbank database (accessed December 2019) (35) to identify the sequences.

### Detailed characterization of 3 hydrogen-sulfide producing isolates

We selected 3 isolates that could aerobically produce H_2_S for further detailed characterization. We selected these isolates because some of the 18 isolates that produced H_2_S when grown with cysteine had identical 16S rRNA sequences, therefore we chose isolates that had distinct 16S rRNA sequences for full-genome sequencing. Additionally, using 16S rRNA gene sequencing of the isolates, one was only assigned to *Stenotrophomonas sp.*, and we believed that whole-genome sequencing would enable us to get a higher taxonomic confirmation and more complete information.

We performed DNA extraction using the PowerSoil Powerlyzer kit (Qiagen) without protocol modifications, and sent the genomic DNA for full genome sequencing at the Microbial Genome Sequencing Center (MIGS) (Pittsburg, PA) with combined short read Illumina and long read nanopore sequencing. The data was processed by MIGS to assemble the short-reads (Illumina Next Seq 2000) and long-reads (Oxford Nanopore Technologies) into full-genomes. Quality control and adapter trimming was performed with bcl2fastq (Illumina) and porechop (https://github.com/rrwick/Porechop) for Illumina and Oxford Nanopore Technologies (ONT) MinION sequencing respectively. Hybrid assembly with Illumina and ONT reads was performed with Unicycler (36). Genome annotation of the 3 isolates was done with Prokka v.1.14.5(37), using the --rfam setting.

Genome completeness and contamination were estimated using CheckM v.1.1.3 (38) *lineage_wf*. Taxonomic classification was conducted using GTDB-tk v.0.3.2 (39) with the database release r95. The full-genome taxonomic classification agreed with the prior 16S rRNA sequencing results, but we were further able to identify Isolate 43 as *Stenotrophomonas bentonitica*. We ran METABOLIC-G v.4.0 (40) to identify genes associated with cysteine degradation and other metabolic pathways.

Growth measurements of the three isolates were measured using OD600 with a spectrophotometer, with measurements every 1 hour. The isolates were grown in R2A broth media, shaken in an incubator at 27°C. Aliquots were collected over the growth range for cysteine measurements across the growth curves, as described below. A H_2_S microsensor (Unisense) was used to measure H_2_S over time.

### Methods to measure Cysteine

Cysteine concentrations were measured as cystine, as described in (41) (https://osf.io/9k8a6/). One of the reasons for measuring cystine instead of cysteine is that in oxic environments, cysteine is oxidized rapidly into cystine (14, 15). Additionally, unless LC-MS is used, cysteine can be difficult to measure directly. Samples were diluted 5:4:1 Sample:DI H_2_O:DMSO and left at room temperature for at least 24 hours. Chromatographic analysis was performed on an Agilent 1260 Infinity II with an Agilent Zorbax Eclipse Plus C18 RRHT 4.6×50mm, 1.8µm, with Guard column. Column temperature was maintained at 40°C using an Agilent 1260 TCC (G1316A).

Gradient elution was performed using Mobile phase A (MPA) consisting of 10mM Na_2_HPO_4_, 10mM Sodium tetraborate decahydrate, in DI H_2_O, adjusted to pH 8.2 with HCl, filtered to 0.45µm. Mobile phase B (MPB) consisted of 45:45:10 Acetonitrile:Methanol:DI H_2_O. Gradient used for elution was as follows: 0 minutes 98% MPA, 2% MPB; 0.2 minutes 98% MPA, 2% MPB; 6.67 minutes 46% MPA, 54% MPB; 6.77 minutes 0% MPA, 100% MPB; 7.3 minutes 0% MPA, 100% MPB; 7.4 minutes 98% MPA, 2% MPB; 8 minutes 98% MPA, 2% MPB. Flow rate was 2.0mL/min. The pump used was an Agilent Infinity Series G1311B quat pump. Pre-column derivatization was performed using an Agilent 1260 ALS (G1329B) with an injector program. Detection was performed using an Agilent 1260 Infinity II MWD (G7165A) at 338nm with 10nm bandwidth. Reference was 390nm with 20nm bandwidth. Recovery was tested during method development. Recoveries of cystine ranged from 87.2-101.5%, with an average of 92.1%.

### Methods to measure H_2_S using a microsensor

Aliquots of at least 1 mL were taken from cultures at desired times after inoculation. H_2_S concentrations were measured by suspending the H_2_S probe (Unisense) in the aliquot and leaving it in place until the measurement stabilized. Measurements were manually edited to exclude data gathered while the probe was stabilizing in the sample.

### Generation of Metagenome-assembled genomes (MAGs)

Sequencing of the Lake Mendota time series for 2008-2012 was previously conducted at the Joint Genome Institute (42), containing 97 time points (and therefore 97 metagenomic datasets)(43). In summary, raw reads were quality filtered using fastp (44), and individually assembled using metaSPAdes(45). Each metagenome was reciprocally mapped to each individual assembly using BBMap v38.07 (46) with 95% sequence identity cutoff. Differential coverage mapping to all samples was used to bin contigs into metagenome-assembled genomes (MAGs) using Metabat2 v.2.12.1 (47). Bins were quality assessed with CheckM v.1.1.2 (38), dereplicated with dRep v.2.4.2 (48), and classified with GTDB-tk v.0.3.2 (49) with default settings. This resulted in a total of 116 MAGs from Lake Mendota (**Table S12**) which are available for download at the Open Science Framework (https://osf.io/qkt9m/).

### Searching for cysteine genes and isolates presence in metagenomic time-series and MAGs

Genes for cysteine degradation were identified using HMMsearch v3.1b2 (50). HMMs were downloaded from KoFam (51), accessed May 2020). The KO numbers for the six cysteine degradation genes are: *metC* (K01760), *cysK* (K01738), *cysM* (K12339), *malY* (K14155), *tnaA* (K01667), and *dcyD* (K05396) (**Table S4**). Our HMM files are available in **Supplementary File 1,** but are the same as published by KOfam, with the modification of manual addition of the TC thresholds. HMM-based homology searches were conducted on the 97 Lake Mendota metagenomes assemblies as described above.

## Data availability

The 16S rRNA sequences for the 29 H_2_S producing isolates, the full genome sequences (nucleotides and amino acids) for isolates #13, #43 and #66 are available on OSF: https://osf.io/g25eq/ during the peer-review process. The sequences will be deposited in NCBI prior to publication. The 97 metagenomes were previously published in (43) and are available through JGI’s IMG/M and Genome Portal. The MAGs generated in this study can be found on https://osf.io/qkt9m/

## Acknowledgements

We are thankful to Anna Schmidt for collecting the original lake water from Lake Mendota in 2018, and to Adam Breister and Elizabeth Zanetakos who enriched the bacterial isolates during summer 2018. We thank Trina McMahon’s Lab and the Long-Term Ecological Research Network, and the Center for Limnology for their field support and prior work on Lake Mendota. We are thankful for the University of Wisconsin’s Water Science and Engineering Laboratory for the use of their HPLC instrumentation, and James Lazarcik for training and assistance with the instruments. We would like to also thank MIGS (the Microbial Genome Sequencing) center (PA, USA) for their assistance with sequencing and long-read data processing.

This work was supported by the USDA National Institute of Food and Agriculture (NIFA) under grant: Hatch project 1025641. Patricia Tran and Kristopher Kieft received the support from the Anna Grant Birge Memorial Award from the Center for Limnology for support for the project in 2019. Patricia Tran is supported by the Natural Science and Engineering Research Council (NSERC) of Canada Doctoral Fellowship. Samantha Bachand was supported by the National Science Foundation (NSF) Research Experience Undergraduate (REU) Award, and the University of Wisconsin-Madison’ Holstrom Environmental Research Fellowship. Kristopher Kieft was supported by a Wisconsin Distinguished Graduate Fellowship Award from the University of Wisconsin-Madison, and a William H. Peterson Fellowship Award from the Department of Bacteriology, University of Wisconsin-Madison. Elizabeth McDaniel was supported by a fellowship through the Department of Bacteriology at the University of Wisconsin –; Madison. This research was performed in part using the Wisconsin Energy Institute computing cluster, which is supported by the Great Lakes Bioenergy Research Center as part of the U.S. Department of Energy Office of Science. We thank the U.S. Department of Energy Joint Genome Institute for sequencing and assembly (CSP 394) of the Lake Mendota metagenomes.

## Contributions

P.Q.T, S.C.B., J.C.H, K.K, and K.A contributed to study design and conceptualization. P.Q.T, S.C.B, J.C.H and K.K conducted experiments on the isolates. J.C.H performed the chemical analyses of the isolates. P.Q.T, S.C.B and J.C.H conducted genomic analyses. P.Q.T, S.C.B and E.A.M conducted metagenomic analyses. P.Q.T, S.C.B and J.C.H analyzed the data and generated figures. P.Q.T, S.C.B, J.C.H and K.A drafted and edited the manuscript. All authors provided feedback and suggestions.

## List of Supplementary Items

### List of Supplementary Tables

**Table S1.** Description of all initial isolates used in screening

**Table S2.** Quantitative measurement of H_2_S over time in the 3 isolates

**Table S3.** Quantitative measurement of cystine over time in the 3 isolates.

**Table S4.** Description of the 6 genes involved in the degradation of L-cysteine or D-cysteine into pyruvate, ammonia and hydrogen sulfide

**Table S5.** Detailed list of all potential enzymatic activities of genes in the cysteine degradation and biosynthesis pathways.

**Table S6.** METABOLIC table: Full genomic content of the 3 isolates

**Table S7.** HMMsearch table output of 16S rRNA amplicons of the 28 isolates that produce H_2_S from Figure 1 against the Lake Mendota Time Series assembly.

**Table S8.** HMMsearch table output of 16S rRNA amplicons of the 28 isolates that produce H_2_S from Figure 1 against the metagenome-assembled-genomes from the LMTS.

**Table S9.** HMMsearch table output of 16S rRNA amplicons of the 28 isolates that produce H_2_S from Figure 1 against the SSU (16S small ribosomal subunit) of the metagenome-assembled-genomes from the LMTS.

**Table S10.** Cysteine hits the Assembly.

**Table S11.** Cysteine hits to MAGS

**Table S12.** MAG characteristics binned from LMTS.

### List of Supplementary Figures

**Figure S1.** H_2_S and ammonia production screening of the isolates

### List of Supplementary Files

**File S1.** HMM profiles.zip

## Supplementary Figures

**Figure S1.**
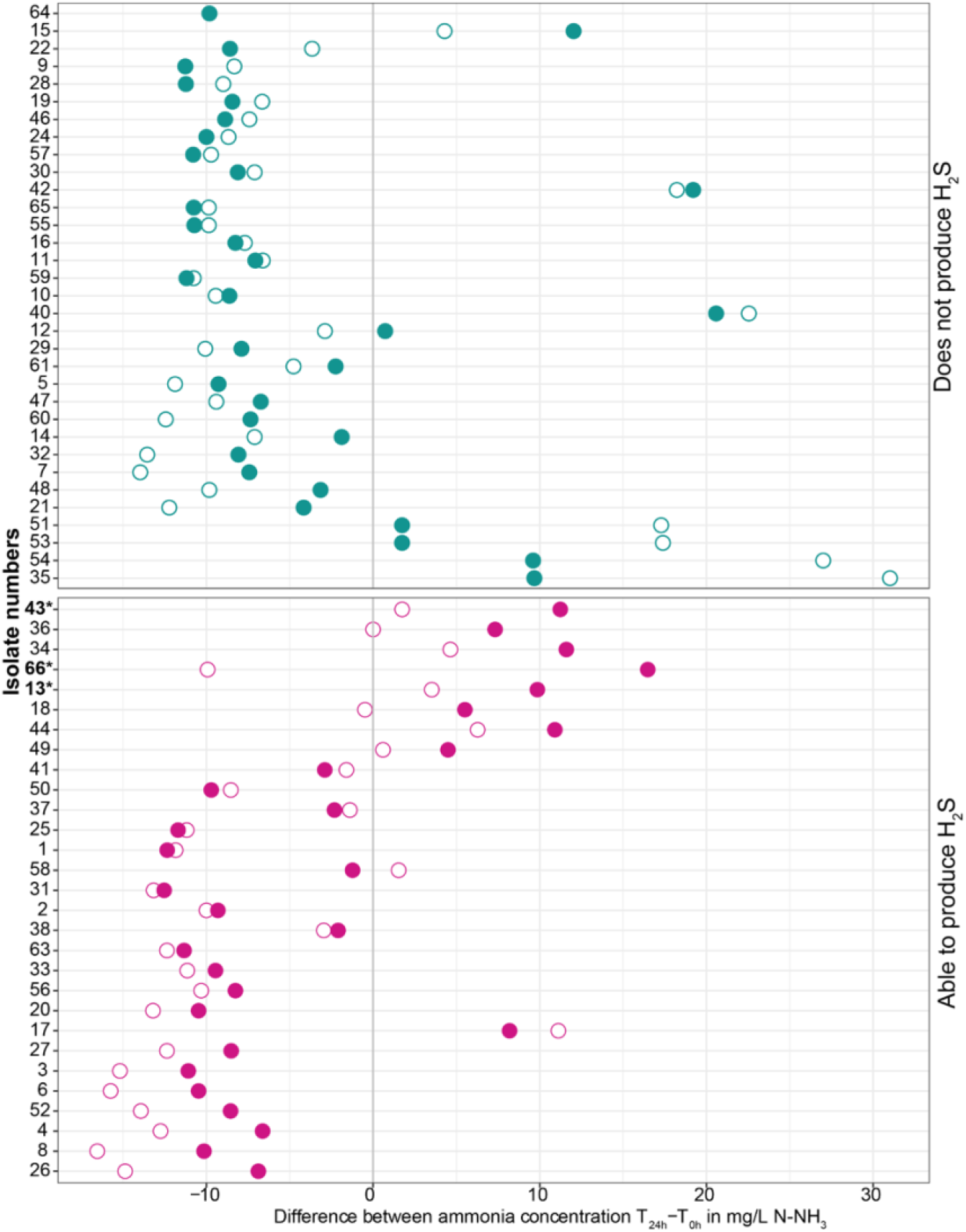
Qualitative accumulation of hydrogen sulfide among microbial isolates enriched from a freshwater lake water column. Filled circles represent isolates grown with cysteine, and open circles represent isolates grown without cysteine. The vertical line represents values corrected for the control (natural ammonia production/consumption in the negative control). All points to the right of the vertical lines indicate an accumulation of ammonia, and all points to the left of the vertical lines refer to those that consumed ammonia after 24 hours. The isolates #43, #13 and #66 (bolded) were selected for further analysis.

## Notes

### Competing Interest Statement

The authors have declared no competing interest.

