## Supplementary figures and images for "Microbial cysteine degradation is a source of hydrogen sulfide in oxic freshwater lakes"

### Supplemental Figure 1

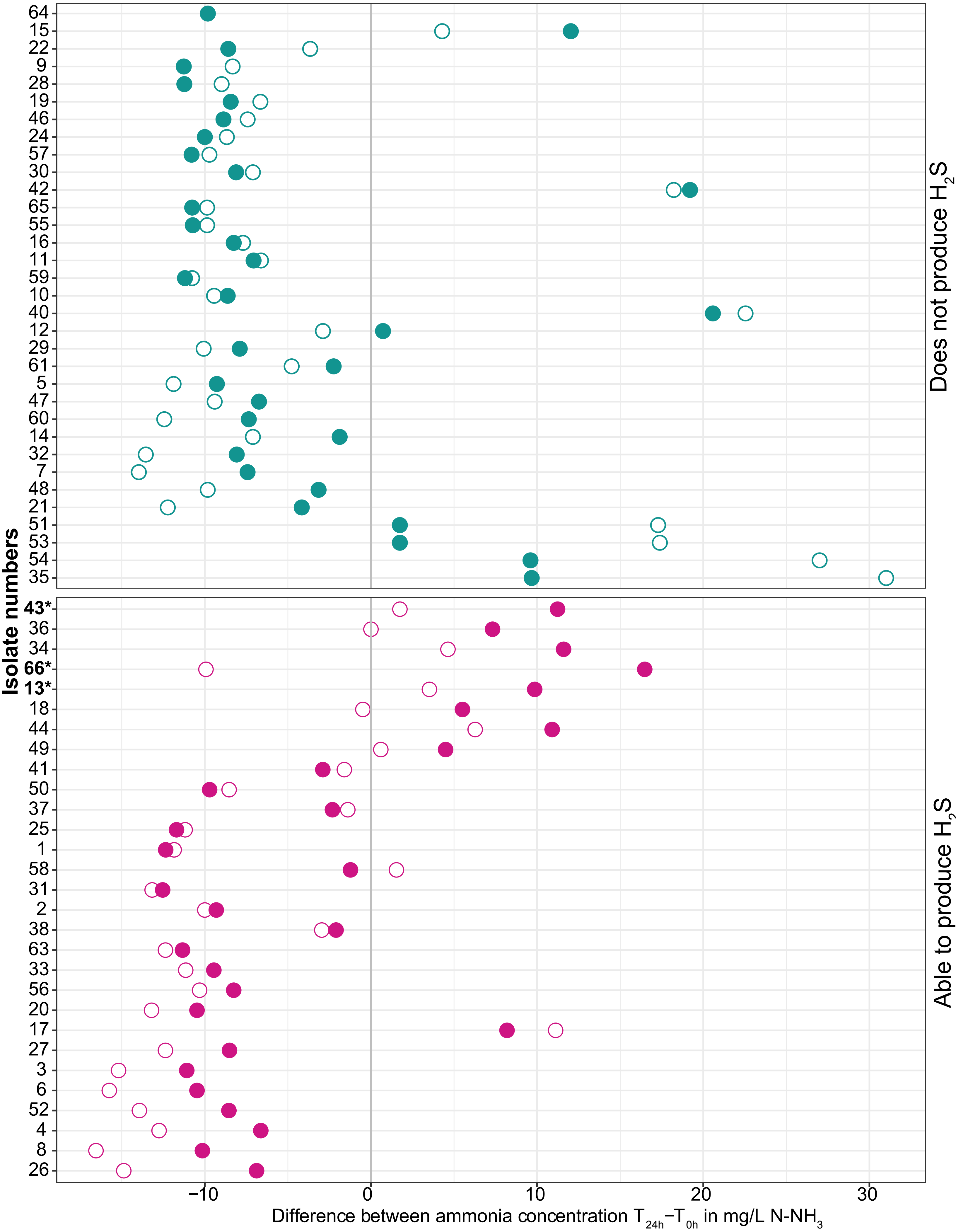
